# Transcriptomic signatures across a critical sedimentation threshold in a major reef-building coral

**DOI:** 10.1101/2023.11.06.565897

**Authors:** Colin Lock, Melissa M. Gabriel, Bastian Bentlage

**Author notes:** Correspondence: PO Box 5067, University of Guam, Mangilao, Guam, USA 96923 –.

## Abstract

Sedimentation is a major cause of global near-shore coral reef decline. While negative impacts of sedimentation on coral reef community composition have been well documented, the effects of sedimentation on coral metabolism *in situ* have received comparatively little attention. Using transcriptomics, we identified gene expression patterns changing across a previously defined sedimentation threshold that was deemed critical due to changes in coral cover and community composition. We identified genes, pathways, and molecular processes associated with this transition that may allow corals, such as *Porites lobata*, to tolerate chronic, severe sedimentation and persist in turbid environments. Alternative energy generation pathways may help *Porites lobata* maintain a persistent stress response to survive while light and oxygen availability are diminished. We found evidence for the expression of genes linked to increased environmental sensing and cellular communication that likely allow *Porites lobata* to efficiently respond to sedimentation stress and associated pathogen challenges. Cell damage increases under stress; consequently, we found apoptosis pathways over-represented under severe sedimentation, a likely consequence of damaged cell removal to maintain colony integrity. The results presented here provide a framework for the response of *Porites lobata* to sedimentation stress under field conditions. Testing this framework and its related hypotheses using multi-omics approaches can further our understanding of metabolic plasticity and acclimation potential of corals to sedimentation and their resilience in turbid reef systems.

## Introduction

Coral reefs fulfill important cultural, ecological, and economic roles in Guam and throughout the Northern Marianas Islands (Burdick *et al*., 2008). Reefs recycle nutrients, provide habitat for marine organisms, and prevent coastline erosion from strong storms and wave action (Hughes *et al*., 2003). Upland erosion caused by human activities such as wild-land arson, deforestation, construction and development, and recreational off-roading have increased turbidity and sedimentation in Guam’s watersheds, exerting significant stress on near-shore coral reef ecosystems (Reynolds *et al*., 2014; Minton, Burdick and Brown, 2022). Wetland and stream destruction by coastal development have further increased sedimentation impacts on near-shore reefs in Guam (Scheman *et al*., 2002; Wolanski *et al*., 2003; Minton, Burdick and Brown, 2022). Increasing erosion of soils and resulting near-shore sedimentation is a wide-spread problem globally, leading to a decline in diversity and ecosystem services provided by coral reefs (Rogers and Ramos-Scharrón, 2022).

Suspended sediment particles increase turbidity and form aggregations that get deposited on corals through sedimentation that negatively impacts coral metabolism (Sheridan *et al*., 2014). Fine sediments and decaying organic matter deposited in nearshore reef systems deplete available oxygen which decreases pH and increases oxygen demand of corals, resulting in oxidative stress (Erftemeijer *et al*., 2012; Flores *et al*., 2012). To survive under such conditions, corals employ mitigation processes, such as increased mucus production and ciliary movement to shed deposited sediment (Weber *et al*., 2012), exerting high energetic costs that can lead to a decrease or cessation of other metabolic functions (Riegl and Branch, 1995; Anthony and Larcombe, 2000). Sedimentation triggers coral immune responses that further deplete energy stores (Sheridan *et al*., 2014) and increase susceptibility to disease (Sheppard, Davy and Pilling, 2009). Increased turbidity caused by suspended sediments cause a decline in photosynthetic activity of coral-associated Symbiodiniaceae (Philipp and Fabricius, 2003; Fabricius, 2005).

Critical thresholds for sedimentation at which coral mortality drastically increases range as low as 10 mg cm^-2^ day^-1^ to as high as 300 mg cm^-2^ day^-1^, depending on coral species impacted, reef location, types of sediments deposited, and length of exposure to sedimentation (Erftemeijer *et al*., 2012; Tuttle and Donahue, 2022). Massive *Porites* spp., such as *Porites lobata*, are one of the few coral species groups known for their persistence under moderate to severe sedimentation (Rogers, 1990; Golbuu *et al*., 2011). Documented responses of *Porites lobata* to sedimentation range from mortality caused by sedimentation rates as low as 30 mg cm^-2^ day^-1^ (Hodgson, 1990) to bleaching but no apparent mortality under severe sedimentation of 200 mg cm^-2^ day^-1^ for 6-8 days (Stafford-Smith, 1993). Large discrepancies in reported sedimentation tolerances of *Porites lobata* may suggest physiological plasticity within this species or represent uncertainties in identification of massive *Porites* spp. and species-specific differences in acclimation potential. Regardless, the persistence of massive *Porites* spp. in environments affected by moderate to severe sedimentation represents unique opportunities for identifying mechanisms of acclimation to sedimentation.

Transcriptome-level characterization of gene expression in corals allows for insights into complex metabolic processes which furthers our understanding of how corals respond to stressors, such as sedimentation, and ultimately, their future survival (Barshis *et al*., 2013, 2014). This approach has been used successfully to lay the foundation for identifying the molecular mechanisms that allow certain coral species to survive or thrive in marginal habitats (Barshis *et al*., 2014). Understanding thresholds of sedimentation tolerance in corals may inform decision-making processes for coral reef conservation and restoration (Burdick *et al*., 2008; Barshis *et al*., 2014; Hughes *et al*., 2017; Tuttle and Donahue, 2022). Studying gene expression of resilient *Porites* corals *in situ* may offer insights into their responses to environmental stressors and adaptation strategies (Barshis *et al*., 2013). Consequently, we employed transcriptomics to characterize the gene expression profiles of *Porites lobata* colonies tagged *in situ* for repeat sampling (Figs. 1a & b) to understand the gene expression profiles of *Porites lobata* and its endosymbiotic Symbiodiniaceae community growing in a habitat characterized by moderate to severe sedimentation. Our analyses identified significant changes in gene expression profiles related to energy metabolism, immune responses, and apoptosis across a previously identified critical threshold of sedimentation across which coral mortality increases and community changes significantly in Fouha Bay, southern Guam (Minton, Burdick and Brown, 2022). Our results obtained from corals growing under natural field conditions are in agreement with prior experimental work conducted under laboratory conditions (Bollati *et al*., 2021), suggesting that the previously proposed acclimation mechanisms to turbid, sedimentation-impacted reef ecosystems are acting on an ecologically relevant scale.

**Figure 1.**
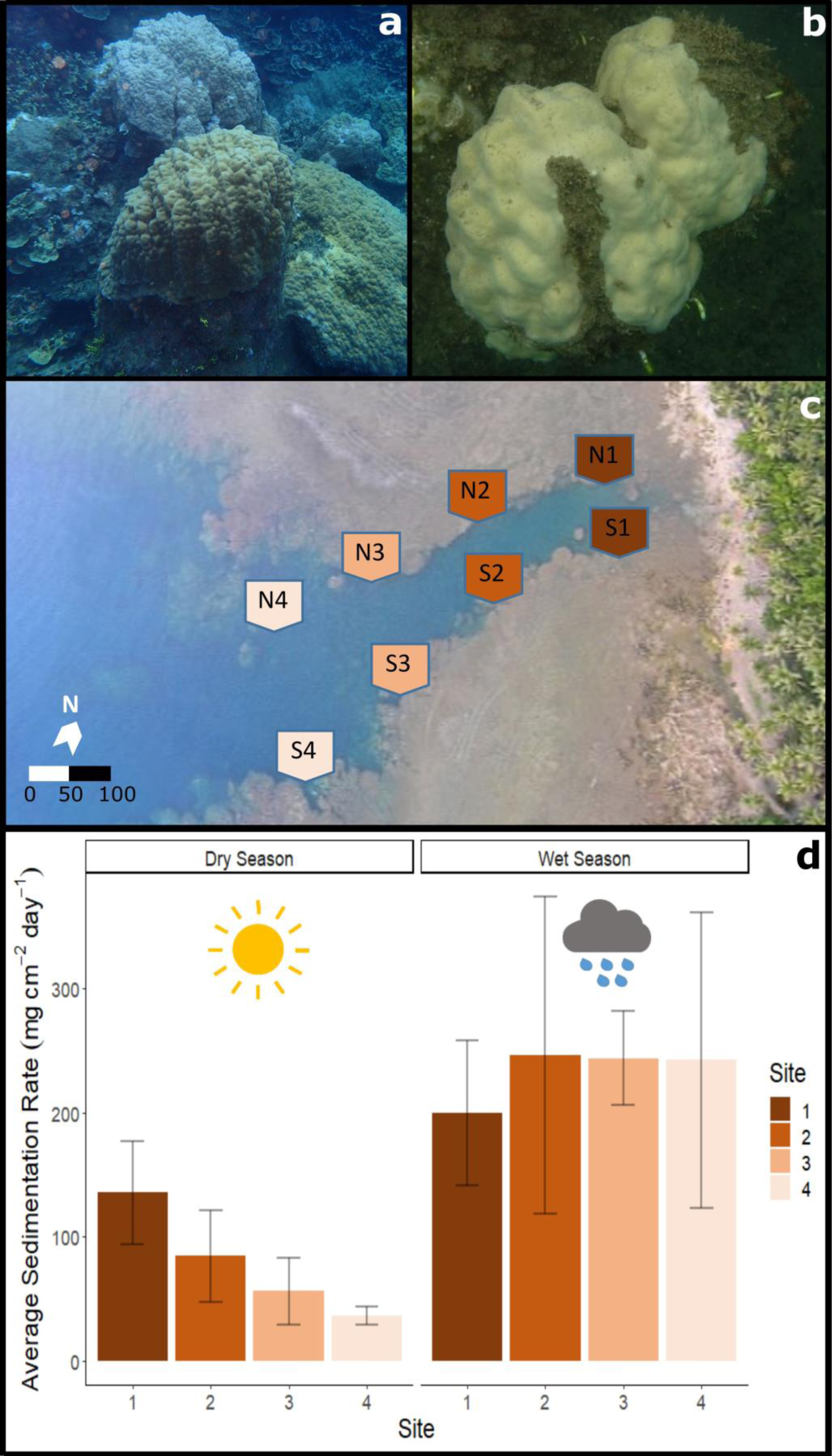
Fouha Bay (Guam) sampling locations and sedimentation rates. **a:** *Porites lobata* colony sampled from the outer zone (N4) that shows minimal sediment accumulation. **b:** *Porites lobata* colony sampled from the inner location (S1) that shows heavy sediment accumulation. **c:** Aerial view of Fouha Bay and sampling sites. **d:** Average sedimentation rate across sampling sites (N/S 1-4) during the dry and wet seasons.

## Material and methods

### Study site and environment

The Humåtak watershed in southern Guam (13°28’ N, 144°45’ E) is characterized by basaltic rock, steep slopes, and lateritic soils that readily erode during heavy rain falls (Foster and Ballendorf, 2023). The La Sa Fu’a river and its 5 km^2^ catchment area belong to the Humåtak watershed that drains into Fouha Bay (Scheman *et al*., 2002), providing mean monthly freshwater discharges ranging from 0.3 m^3^ s^-1^ (March – May) to 2.9 m^3^ s^-1^ (August – October) (United States Geological Survey monitoring site 16809600 data averaged from 1953 to 2018; Geological Survey, no date), corresponding to dry and wet season rainfall patterns. Heavy rain events that occur primarily during the wet season and last on average one to two hours, transport sediment from eroding soils downstream into Fouha Bay, creating pulses of thick plumes that frequently exceed a suspended solids concentration (SSC) of 1000 mg l^-1^ (Wolanski *et al*., 2003). Suspended solids flocculate and slowly precipitate as marine snow when reaching the bay, dissipating the plume over a time-span of some five days (Wolanski *et al*., 2003). Large sedimentation events occur on average ten times annually in Fouha Bay with some 75% of sediments staying within the bay for an estimated residence time of 4.5 years (Wolanski *et al*., 2003).

Massive *Porites* of the *Porites lobata*/*lutea* species complex (Figs. 1a & b) represent one of the few coral species groups that persist in Fouha Bay along a gradient of sedimentation ranging from severe (sedimentation rate >50 mg cm^-2^ day^-1^) to moderate (sedimentation rate ∼10 – ∼50 mg cm^-2^ day^-1^) to light (sedimentation rate <10 mg cm^-2^ day^-1^), as distance from the river mouth increases (Minton, Burdick and Brown, 2022). Sites close to the river mouth are further characterized by decreased light available for coral photosynthesis due to high turbidity and decreased salinity due to freshwater discharge (Fifer *et al*., 2022).

To evaluate previously established sedimentation dynamics across Fouha Bay (Rongo, 2004; Minton, Burdick and Brown, 2022), sedimentation rates during the period of this study were estimated using a single sediment trap deployed at 8 sites established within the bay following Rongo (2004) (Fig. 1c). Sediment was collected over a time period of 31 days during the dry (May 2, 2018 to June 1, 2018) and wet (September 21, 2018 to October 22, 2018) seasons using vertical PVC traps (5 cm diameter and 50 cm height), following the recommendations of (Storlazzi, Field and Bothner, 2011). Processing of collected sediments followed the protocol of (Ensminger, 2016). Wet-strengthened 120-micron filters (Whatman, Little Chalfont, UK) were first rinsed with deionized water and dried for 24 hours at 100° C; weight of each dried filter was recorded. Collected sediment samples were rinsed with deionized water to remove salt and allowed to settle again for 24 hours; organisms contained in sediment samples were removed. Cleaned samples were then filtered through previously dried 120-micron filters using a vacuum pump, followed by drying of sediments collected on filter paper at 100° C. Drying filters and sediments were weighed repeatedly over a period of two days until weights remained steady, indicating that samples had dried completely, allowing for calculation of accumulated sediment. Normalization of accumulated sediment to deployment time and diameter of collection tube yielded sedimentation rate in mg cm^-2^ day^-1^. A Shapiro-Wilks test was used to test normality in the sedimentation dataset and then an ANOVA was used to compare sedimentation rates between sites, seasons, and sides (using the AOV function in base R; version 4.2.3).

### Coral tagging and identification

At each of the eight sites in Fouha Bay (Fig. 1c), at least three corals were tentatively identified in the field as *Porites lobata* based on gross morphology and tagged for repeated sampling. Coral colonies ranged in diameter from roughly 15 cm to 2 m, with colonies closer to the river mouth being smaller than more distant ones. Species were identified using a combination of corallite morphology and, given the difficulty of species-level identification of massive *Porites* species, DNA barcoding. Fragments from all 24 tagged colonies were collected with hammer/chisel and preserved in ethanol for DNA extraction.

DNA was extracted from each tagged colony using the GenCatch genomic DNA extraction kit (Epoch Life Science, Sugar Land, TX) following the manufacturer’s protocol for tissue samples. Mitochondrial regions COX3-COX2 and ND5-tRNA-Trp-ATP8-COX1 were amplified with primer sets mt-16 and mt-20 (Paz-García *et al*., 2016) in 25 μl reactions using 0.3uM primer, 0.3mM dNTP, 1x HiFi Fidelity Buffer, and 2.5 units Taq (HIFI kapa 1U). The thermocycler profile included an initial denaturation at 94°C for 120 s, followed by 30 cycles of 94°C for 30 s, 54°C for 30 s, and 72°C for 60 s, followed by a final extension at 72°C for 300 s. PCR products were sequenced using Sanger sequencing and resulting sequences assembled using the overlap-layout-consensus algorithm implemented in Geneious Prime (Biomatters, Auckland, New Zealand). Following assembly, consensus sequences for each specimen were aligned to publicly available *Porites* spp. mitochondrial genomes using MAFFT v7.453. The maximum likelihood phylogeny was inferred from the concatenated, aligned regions using RAxML v8.2.12 (Stamatakis, 2014) under the GTRCAT model to allow for efficient modeling of site heterogeneity across alignment regions that spanned multiple genes. The resulting phylogeny was rooted using *Porites fontanesii* following recent phylogenomic analyses (Terraneo *et al*., 2021); robustness of the phylogeny was assessed using 1,000 nonparametric bootstrap replicates. Specimens grouping with *Porites lutea* were excluded from RNA sequencing while specimens grouping with *Porites lobata* were included in the set of specimens selected for gene expression analysis.

### RNA extraction, sequencing, and transcriptome assembly

Tagged colonies were sampled during the dry season on May 16, 2018 and on October 3, 2018 during the wet season, following the passing of two tropical depressions that lead to significant rainfalls. A fragment of each coral colony was sampled using hammer/chisel and immediately preserved in RNA later (Sigma-Aldrich, St. Louis, MO, USA), followed by storage at −80°C until RNA extraction. RNA from specimens identified as *Porites lobata* (see section above) was extracted using the E.Z.N.A Mollusc RNA kit (Omega Bio-tek, Norcross, GA USA), following the manufacturer’s protocol. Extracted RNA was quantified using a Qubit fluorometer (Life Technologies, Carlsbad, CA) and its integrity verified using a Bioanalyzer (Agilent Technologies, Santa Clara, CA), leading to the selection of 32 samples for RNA sequencing. Sequencing libraries were constructed using NEBNext library kits (New England Biolabs, Ipswich, MA, USA) and sequenced on a NextSeq 550 sequencer (Illumina, San Diego, CA, USA), generating 150bp paired-end data. Adapter sequences and bases with a phred-scaled quality score of less than 30 were removed using TrimGalore (Martin, 2011). Sequence data were deposited in NCBI’s GenBank (accession numbers provided in Table S1).

We employed *de novo* transcriptome assembly following best practices outlined in Grabherr et al. (2013). After exclusion of two samples that yielded few sequencing reads (Table S2), sequencing data were combined and normalized using the Trinity v2.10.0 (Grabherr *et al*., 2013) *in silico* normalization function with maximum coverage set to 50 to reduce the computational requirements of transcriptome assembly that used Trinity’s default parameters. After transcriptome assembly, TransDecoder v5.5.0 (Haas *et al*., 2013) was used to predict open reading frames (ORFs), followed by removal of non-coding transcripts. Non-target contaminant sequences were identified and removed from the transcriptome assembly using the PERL script Alien Index (Ryan, 2014) with the approach outlined by (Lock, Bentlage and Raymundo, 2022). Representative proteomes inferred from translated coding sequences (391,427 sequences) for each major bacterial clade (Schulz *et al*., 2017), fungi, stramenopiles, poriferans, arthropods, molluscs, and annelids were obtained from NCBI’s GenBank and indexed in a non-target (alien) BLAST (Altschul *et al*., 1990) database; 57,401 coral and 72,664 Symbiodiniaceae proteome sequences were included as target sequences (File S1). Protein BLAST searches (e-value < 1e^-3^) using the open reading frames (ORFs) predicted from our transcriptome assembly by TransDecoder as queries were run against the combined non-target (alien) and target (Cnidaria plus Symbiodiniaceae) protein database to identify and remove likely contaminant sequences from our *de novo* assembled transcriptome with Alien Index (Ryan, 2014). Target coral and Symbiodiniaceae ORFs were annotated based on the best BLAST hit (e-value < 1e-5) against a database containing 129,600 cnidarian (including corals and other cnidarians) and 44,114 Symbiodiniaceae amino acid (protein) sequences (Table S3) obtained from the UniProt database (Bateman *et al*., 2017).

Benchmarking Universal Single Copy Orthologs (BUSCO) v5.1.2 (Simão *et al*., 2015) was used to estimate the completeness of the taxonomically filtered *Porites lobata* and Symbiodiniaceae transcriptomes. The transcriptome assemblies for *P. lobata* and Symbiodiniaceae were compared against the metazoan and alveolate BUSCO gene sets, respectively. For *de novo* assembled transcriptomes, it is expected that multiple predicted isoforms map to the same BUSCO gene, which can lead to high inferred gene duplication rates. To account for this potential artifact, multiple isoforms of the same gene model that mapped to the same BUSCO gene were counted as a single hit.

### Identification of Symbiodiniaceae clades

RNA sequence reads from each sample (Table S2) were mapped against publicly available Symbiodiniaceae transcriptomes (Bayer *et al*., 2012; Ladner, Barshis and Palumbi, 2012) using a custom Perl script (zooxType3.pl; Manzello et al., 2019). The number of high-quality mapped reads (MAPQ > 40) to representative Symbiodiniaceae transcriptome assemblies was used to calculate the proportions of *Symbiodinium, Breviolum, Cladocopium,* and *Durusdinium* in the sampled *Porites lobata* colonies (Manzello *et al*., 2019).

### Differential gene expression and GO enrichment

Transcriptomes assembled *de novo* usually generate many more gene models and transcripts than predicted from whole genome sequences with read coverage frequently not being uniform across inferred transcripts (cf. Hayer et al., 2015). This is often addressed by collapsing inferred transcripts using clustering based on a similarity cutoff chosen prior to differential gene expression analysis. Rather than reducing the size of our transcriptome through transcript clustering based on an arbitrarily chosen cutoff, we followed the ‘analysis first, aggregation second’ approach of Yi et al. (2018) that addresses the issue of uneven read coverage across transcripts by aggregating *p* values from transcript-level differential expression analysis to identify differentially expressed genes (DEGs). Transcript abundances were estimated using the fast k-mer hashing and pseudoalignment algorithm implemented in Kallisto v0.46.2 (Pimentel *et al*., 2017), followed by differential gene expression analysis in R using the Sleuth v0.30.0 (Pimentel *et al*., 2017) package. Read counts for transcripts belonging to the same gene were aggregated (p-value aggregation;(Pimentel *et al*., 2017)) to infer differential expression at the gene level. A likelihood ratio test comparing a full and reduced model was used to identify significant DEGs between sites, locations, and seasons while accounting for variation of other factors incorporated in the two models (full model: Site + Season + Side [N or S]; null model: Season + Side). All pairwise comparisons between sites (1 vs 2, 2 vs 3, etc.) were assessed for DEGs (*q*-value < 0.05) with side of Fouha Bay (N or S) and season used for the null model.

Since comparisons 1 vs 2 and 3 vs 4 showed extremely low DEGs, sites 1 and 2 were combined into Inner (closer to the river), sites 3 and 4 into Outer (closer to the opening of Fouha Bay) for further comparisons. Wald’s tests were used within sleuth to obtain log2 fold changes of DEGs between conditions. Genes identified as differentially expressed were separated into up and down regulated (enriched in the Inner and Outer zone, respectively; Fig. 4) and used for GO enrichment analysis using the R package GO_MWU using default parameters (Wright *et al*., 2015). The R packages pheatmap (Version 1.0.12) and ggplot2 (Version 3.4.2) were used to visualize the heatmap and GO enrichment plots.

## Results

### Seasonal sedimentation patterns

During the dry season, sedimentation rates followed previously measured and modeled sedimentation rates for Fouha Bay (Rongo, 2004; Minton, Burdick and Brown, 2022), showing a gradient from severe sedimentation, exceeding rates of 135 mg cm^-2^ day^-1^ near the river mouth at sites N1 and S1, to moderate rates of 36 mg cm^-2^ day^-1^ at sites N4 and S4 (Figs. 1c & d).

Comparison of the inner (Sites 1 & 2) and outer (Sites 3 & 4) zones showed statistically significant differences in sedimentation rates for the dry season (p = 0.029). During the wet season, sedimentation rates converged, being extremely severe across all sites (p = 0.993; Fig. 1d). Average sedimentation rates did not differ between seasons for sites N1 and S1, but sedimentation increased for the remaining sites during the wet season compared to the dry season (Fig. 1d). These seasonal differences in sedimentation rates were mirrored by differences in rainfall with precipitation recorded at the Humåtak rain gauge being almost twice as high during the wet season from September 21, 2018 to October 22, 2018 (∼15 mm day^-1^) compared to the dry season from May 2, 2018 to June 1, 2018 (∼8 mm day^-1^) (US Geological Survey monitoring site 131729144393766; Geological Survey, no date).

### Species identification, sample selection, and RNA sequencing

Massive *Porites* species are morphologically cryptic (Forsman *et al*., 2015). Using a DNA barcoding approach, we assigned 20 tagged coral colonies to *Porites lobata*, forming a well-supported monophyletic clade, while 4 colonies were assigned to *Porites lutea* (Fig. 2a) and excluded from further analyses. RNA was extracted from the 20 coral colonies identified as *Porites lobata.* For each site (N1 to N4 and S1 to S4; Fig. 1c), samples that produced the best quality RNA extracts (RIN > 6) were chosen for sequencing, yielding a total of 30 samples (Tables S1 & S2; Fig. 1c) for differential gene expression analysis.

**Figure 2.**
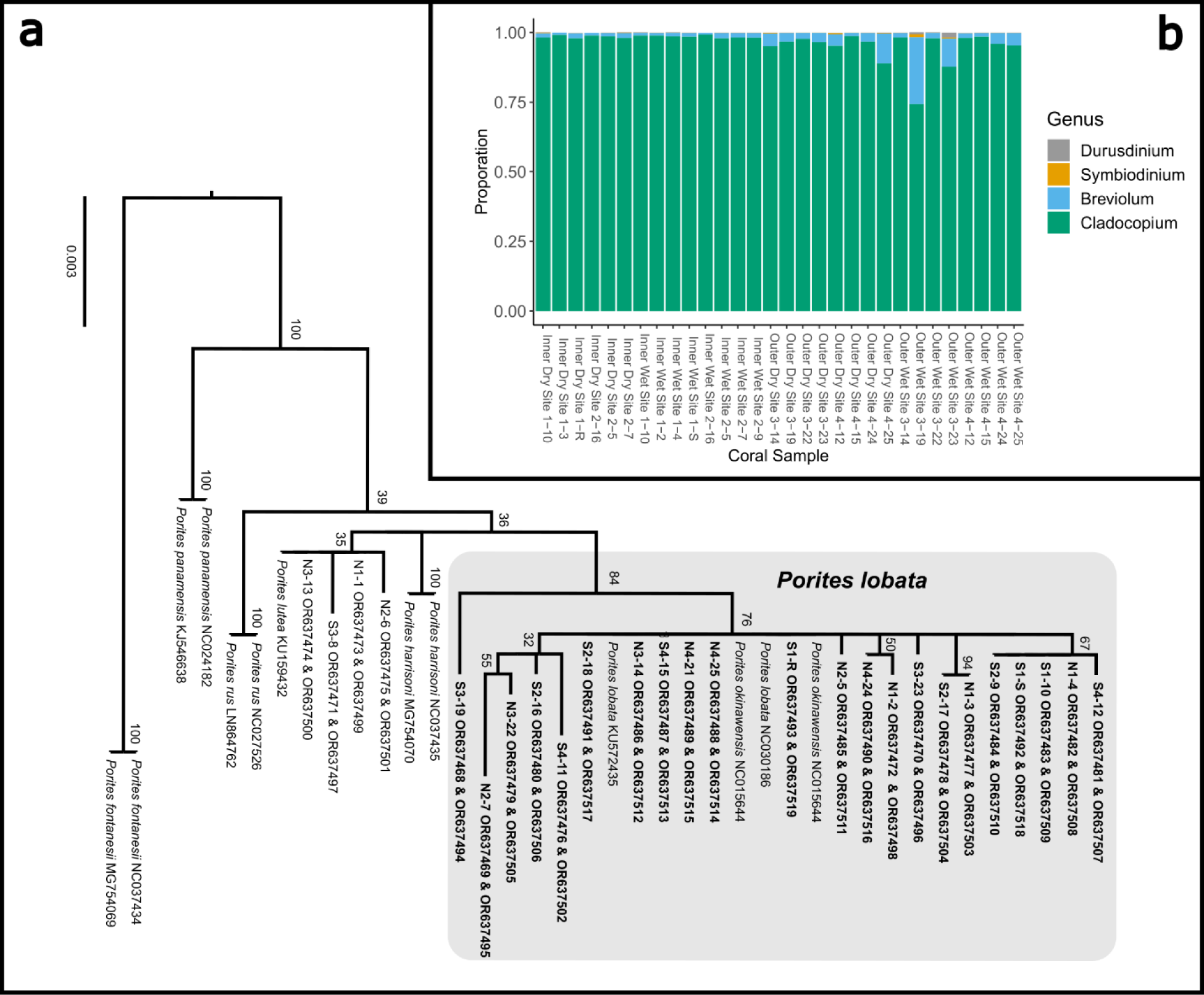
**a:** Maximum likelihood phylogeny of concatenated mitochondrial COX3-COX2 and ND5-tRNA-Trp-ATP8-COX1 inferred under the GTRCAT model implemented in RAxML (Stamatakis, 2014). Node support was evaluated using 1,000 non-parametric bootstrap replicates. Tagged *Porites* colonies indicated by site, followed by specimen number (e.g., N2-6 represents specimen tag 6 from site N2); colonies in bold were sampled for transcriptome sequencing. ORXXXXXX are the GeneBank accession numbers for that sample **b:** Proportionate contributions of Symbiodiniaceae clades contained in each *P. lobata* sample based on read mapping to reference transcriptomes.

RNA sequencing generated 81,379,001 ± 55,399,881 (mean ± SD) reads with 73,696,495 ± 51,328,889 reads retained after quality trimming. After *de novo* transcriptome assembly of the filtered reads, removal of contaminants, and taxonomic binning, the resulting *Porites lobata* and Symbiodiniaceae transcriptomes contained 72,813 putative gene models (105,623 transcripts) and 57,891 putative gene models (72,428 transcripts), respectively. The transcriptome of *Porites lobata* was relatively complete with almost 90% of BUSCO genes complete (C:88.5% [S:78.2%, D:10.3%], F:5.0%, M:6.5%, n:954); the Symbiodiniaceae transcriptome contained close to 70% of complete BUSCO genes (C:67.8% [S:44.4%, D:23.4%], F:6.4%, M:25.8%, n:171). Relatively high levels of BUSCO gene duplication are expected in transcriptome assemblies compared to genome assemblies, as multiple transcripts (isoforms) may map to the same BUSCO gene.

Mapping of Symbiodiniaceae transcripts to publicly available Symbiodiniaceae transcriptomes revealed that all *Porites lobata* colonies predominately harbored (> 74% relative abundance) *Cladocopium* spp., with most samples showing > 95% abundance of *Cladocopium* spp. (Fig. 2b; Table S4).

### Differential gene expression and GO term enrichment

Pairwise comparisons between sites produced similar results for *Porites lobata* and Symbiodiniaceae: fewer than two genes were differentially expressed when comparing between sites 1 and 2 as well as 3 and 4 (Table 1). Given these results, sites were combined into inner zone (sites 1 and 2) and outer zone (sites 3 and 4) that correspond to the transition between moderate and severe sedimentation. Taking variation between zones (inner *versus* outer) and side (N or S) into account, season produced 5 differentially expressed genes (DEGs) for *Porites lobata* and none for Symbiodiniaceae (Table 2). Additionally, comparisons of wet *versus* dry season for only the inner zone and only the outer zone revealed between 0 and 11 DEGs for *Porites lobata* and none for the Symbiodiniaceae (Table 2). Side of the Fouha Bay (North *versus* South) produced 42 differentially expressed genes for *Porites lobata* and none for Symbiodiniaceae.

**Table 1.** Number of differentially expressed genes for each pairwise comparison between sites using likelihood ratio model comparisons; factors included in null models given in brackets. Sites were merged into inner (sites 1 & 2) and outer (sites 3 & 4) zones based on the low gene expression between those sites.

| *Porites lobata*: inner *versus* outer zone<br>[side (N/S) + season] |  |  |  |  | Symbiodiniaceae: inner *versus* outer zone<br>[side (N/S) + season] |  |  |  |  |
| --- | --- | --- | --- | --- | --- | --- | --- | --- | --- |
| Site | 1 | 2 | 3 | 4 | Site | 1 | 2 | 3 | 4 |
| 1 | - |  |  |  | 1 | - |  |  |  |
| 2 | 2 | - |  |  | 2 | 0 | - |  |  |
| 3 | 736 | 499 | - |  | 3 | 825 | 97 | - |  |
| 4 | 873 | 1,333 | 0 | - | 4 | 435 | 488 | 0 | - |

**Table 2.** Number of differentially expressed genes identified for *Porites lobata* and its Symbiodiniaceae endosymbionts using likelihood ratio model comparisons; factors included in null models given in brackets. The comparison of inner and outer zones yielded the largest number of differentially expressed genes. Arrows indicate the number of genes upregulated (↑) and downregulated (↓) in the inner zone.

| Comparison | *Porites lobata* | Symbiodiniaceae |
| --- | --- | --- |
| Season: dry *versus* wet<br>[side (N/S) + zone] | 5 | 0 |
| Season (inner zone): dry *versus* wet<br>[side (N/S)] | 11 | 0 |
| Season (outer zone): dry *versus* wet<br>[side (N/S)] | 0 | 0 |
| Side: North (N) *versus* South (S)<br>[season + site] | 42 | 0 |
| Zone: inner *versus* outer<br>[season + side (N/S)] | 1,702<br>(↑ 433; ↓ 1,259) | 1,514<br>(↑ 319; ↓ 1,195) |

Zone (inner *versus* outer) produced the largest number of DEGs for both *Porites lobata* (1,702 DEGs; 433 up-regulated and 1,259 down-regulated in the inner zone) and Symbiodiniaceae (1,514 DEGs; 319 up-regulated and 1,195 down-regulated in the inner zone) (Table 1; Table S5). These results mirror patterns of gene expression profile similarity that largely group samples by zone rather than season, differentiating between the inner zone affected by severe sedimentation and outer zone impacted by moderate sedimentation (Fig. 3). GO enrichment analysis (Table S7) identified 38 biological processes enriched (p < 0.01) in *Porites lobata* (Fig. 4a & c) and 10 biological processes enriched (p < 0.01) in Symbiodiniaceae (Fig. 4b & d), which were separated into GO categories enriched in the inner and outer zone.

**Figure 3.**
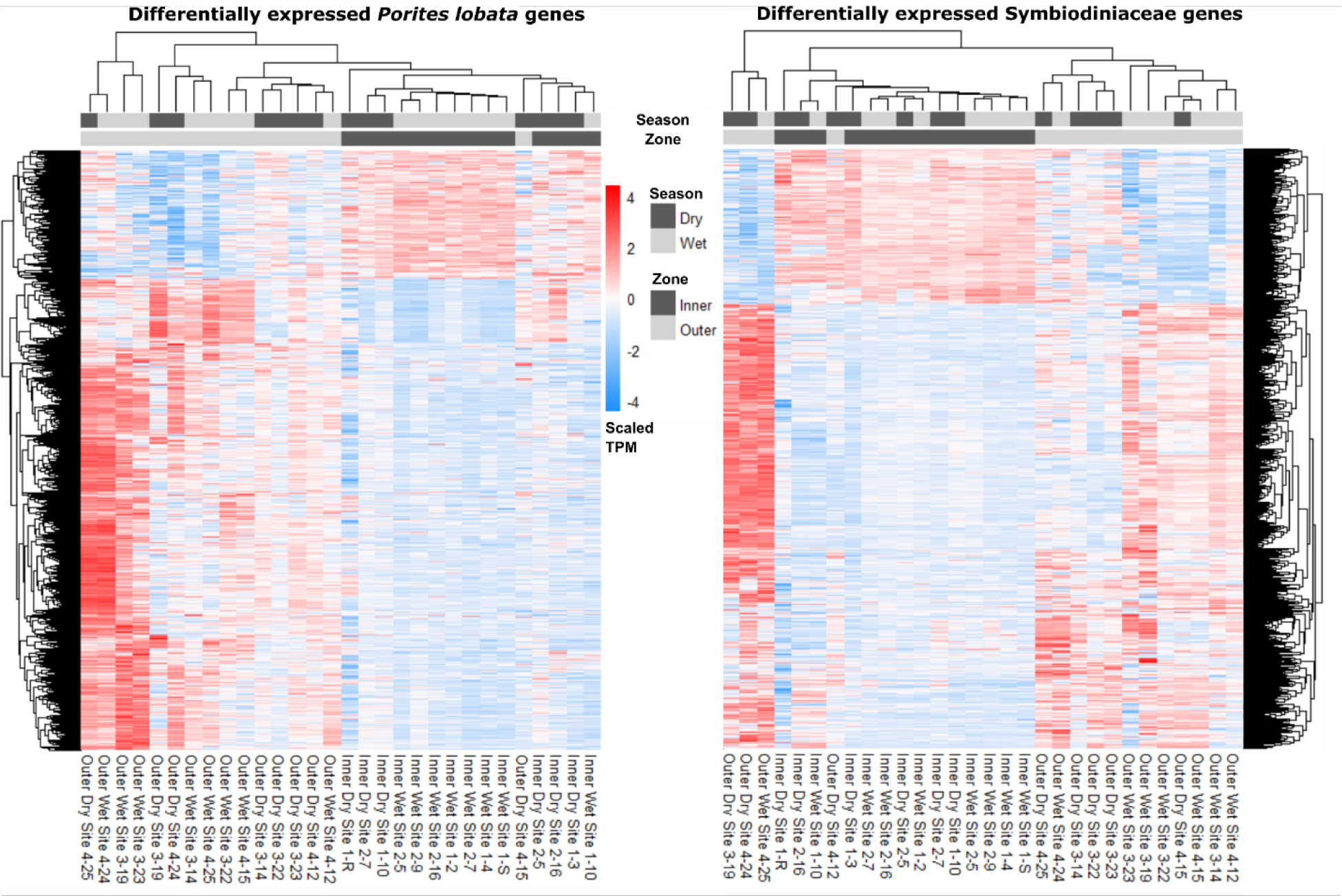
Heatmaps depicting read counts in transcripts per million (TPM) for each differentially expressed gene (rows) for *Porites lobata* and Symbiodiniaceae. Samples (columns) are ordered by gene expression similarity. For both *Porites lobata* and Symbiodiniaceae, samples cluster by zone rather than season (see results for more information).

## Discussion

During our sampling periods, sedimentation ranged from moderate (37 mg cm^-2^ day^-1^) to severe (136 mg cm^-2^ day^-1^) during the dry season along the environmental gradient sampled in Fouha Bay (Fig. 1d). During the wet season, all sites were inundated with extremely severe levels of sedimentation (>200 mg cm^-2^ day^-1^; range: 199-245 mg cm^-2^ day^-1^) (Fig. 1d). Our observations are congruent with previous studies that documented and modeled sedimentation rates at times exceeding 200 mg cm^-2^ day^-1^ in Fohua Bay (Randall and Birkeland, 1978; Wolanski *et al*., 2003; Rongo, 2004; Minton, Burdick and Brown, 2022). Rongo’s (2004) model of annualized sedimentation rates (N1/S1 = 102.5 mg cm^-2^ day^-1^, N2/S2 = 65.2 mg cm^-2^ day^-1^, N3/S3 = 53.4 mg cm^-2^ day^-1^, N4/S4 = 22.5 mg cm^-2^ day^-1^) closely mirrors our sampling period in the dry season that shows a gradient of sedimentation rates declining from the river mouth toward the outer parts of Fouha Bay (Fig. 1d), with the transition from moderate to severe sedimentation between inner and outer zones representing a significant difference in sedimentation rates.

However, wet season sedimentation documented by us, albeit with limited sampling, showed that the sedimentation gradient in Fouha Bay broke down during the wet season, with all sites becoming impacted by severe sedimentation. Nonetheless, declines in coral coverage and species richness documented over the last 40 years in Fouha Bay, linked to long-term chronic sedimentation stress, followed the patterns of the dry season sedimentation gradient, with lowest coral diversity and abundance near the river mouth where sedimentation is severe and coral diversity and abundance increasing, as sedimentation drops to moderate or light levels further from the river mouth (Randall and Birkeland, 1978; Minton, Burdick and Brown, 2022). The stark decline of coral diversity and cover in Fouha Bay linked to the transition from moderate to severe sedimentation has previously been identified as a critical sedimentation threshold (Minton, Burdick and Brown, 2022).

Despite a large difference in sedimentation rates between dry and wet seasons (Fig. 1d), gene expression profiles were more similar across samples within zones compared to seasons (Fig. 3; Table 2). Indeed, differential gene expression analysis comparing dry and wet seasons yielded a minimal number of differentially expressed genes in both *Porites lobata* and its Symbiodiniaceae endosymbiont community (Table 2). The lack of detectable gene expression differences between seasons may be linked to the chronic nature of the sedimentation stress experienced by *Porites lobata* in Fouha Bay, consistent with drastic reductions in coral cover and diversity between outer and inner zones (Minton, Burdick and Brown, 2022). The outer zone of Fouha Bay is more exposed to wave action than the inner zone that would allow for more expedient dissipation of turbidity and accumulated sediments. Given that sampling during the wet season occurred following significant rain events, sedimentation rates may have increased temporarily in the outer zone of Fouha Bay, temporarily obscuring the well-described sedimentation gradient of Fouha Bay (Rongo, 2004; Minton, Burdick and Brown, 2022). The lack of differentially expressed genes between sites N3/S3 and N4/S4 in the outer zone and N1/S1 and N2/S2 in the inner zone points to an environmental break between zones, consistent with a transition from moderate to severe sedimentation rates (Fig. 1d) (Minton, Burdick and Brown, 2022). The large number of differentially expressed genes identified when comparing inner and outer zones using differential gene expression analysis (Table 2), in conjunction with clustering of inner and outer zones based on gene expression similarities (Fig. 3), suggests a strong metabolic threshold associated with the transition from moderate (∼57 mg cm^-2^ day^-1^ during the dry season) to severe (∼85 mg cm^-2^ day^-1^ during the dry season) sedimentation in *Porites lobata*. These results are congruent with Minton et al.’s (2022) identification of a critical sedimentation threshold of 48 mg cm^-2^ day^-1^ at which abundance and diversity of corals drastically decline in Fouha Bay, with massive *Porites* spp. being one of the few taxa able to persist.

Taken together, coral community survey data (Minton, Burdick and Brown, 2022) and our gene expression analysis suggest that *Porites lobata* and their Symbiodiniaceae endosymbionts are chronically stressed due to severe sedimentation in Fouha Bay’s inner zone. Through comparisons of gene expression patterns (Fig. 3; Table 2), annotation of differentially expressed genes and GO term enrichment analysis (Tables S3 - S5), we identified the putative signatures of sedimentation on the energy metabolism and immune response of *Porites lobata* and its endosymbiotic Symbiodiniaceae community discussed below. Messenger RNA (mRNA) and protein expression are often poorly correlated (e.g., due to variations in mRNA stability and translational efficiency; (Payne, 2015; Liu, Beyer and Aebersold, 2016) including in corals and their symbionts (e.g., Mayfield et al., 2016). While this has led to concerns about interpreting differential gene expression patterns, the correlation of differentially expressed mRNAs with protein expression may be significantly better than for non-differentially expressed genes (e.g., Koussounadis et al., 2015). Transcriptomics allows for rapidly profiling tens of thousands of potential biomarkers at once which we used to provide a framework of the coral sedimentation stress response under field conditions that can guide multi-omics approaches aimed at linking gene to protein expression and their enzymatic and metabolic products to untangle the regulation and complexity of this system.

### Energy metabolism

GO terms associated with oxidative phosphorylation (GO:0006119) and aerobic respiration (GO:0009060) were enriched for *Porites lobata* in the outer zone, exposed to moderate sedimentation (Fig. 4a). For Symbiodiniaceae, photosynthesis (GO:0015979), generation of precursor metabolites and energy (GO:0006091), electron transport chain (GO:0022900), and lipid transport (GO:0006869) were enriched in the outer zone (Fig. 4c). Photosynthesis-associated genes coding for fucoxanthin-chlorophyll a-c binding proteins A, F, B & E and Photosystem I/II reaction center proteins were downregulated in Symbiodiniaceae from the inner zone (Table S6), suggesting that maintenance of the photosynthetic machinery is reduced under severe sedimentation, likely attributable to depleting energy stores (Sheridan *et al*., 2014), reallocation of energy to crucial stress responses (Riegl and Branch, 1995), reduced light, and/or decreased oxygen availability (Jones and Hoegh-Guldberg, 2001; Weber *et al*., 2012). Other studies have reported significant reduction in maximum quantum yield (F_v_/F_m_) of photosystem II in response to sedimentation stress from a variety of coral species (Philipp and Fabricius, 2003), consistent with decreased rates of photosynthesis and a significant reduction in lipid stores of corals (Sheridan *et al*., 2014). Upregulation of lipid catabolic and transport enzymes (Lipases, Apolipoprotein L domain-containing protein 1; Table S6) and downregulation of fat storage-inducing transmembrane protein 2 (Table S6) in *Porites lobata* indicate that corals in the inner zone of Fouha Bay are likely depleting their lipid reserves to maintain antioxidant production, an innate immune response, and increased cilia action and mucous production to persist under chronic and severe sedimentation stress that requires removal of sediment deposits, mitigation of pathogens, and elimination of increasing reactive oxygen species (ROS) that are the result of decreased dissolved oxygen available for metabolic processes in a turbid environment. Reduction of photosynthesis, as suggested by the downregulation of photosystem genes in Symbiodiniaceae observed in this study may be offset by increased heterotrophic feeding, which has been suggested as an adaptive strategy for corals persisting in turbid environments (Bay *et al*., 2009; Pacherres, Schmidt and Richter, 2013), a possible strategy for *Porites lobata* that is a mixotroph (Conti-Jerpe *et al*., 2020). Taken together, our results of GO term enrichment (Fig. 4) and differential gene expression (Table 2; Table S6) analyses indicate that *Porites lobata* in the inner zone showed a decrease in energy generation, likely due to a depletion of energy reserves (Sheridan *et al*., 2014), and decreased photosynthetic activity of Symbiodiniaceae due to reduced light availability and/or lack of oxygen availability for aerobic respiration (Jones and Hoegh-Guldberg, 2001; Philipp and Fabricius, 2003).

**Figure 4.**
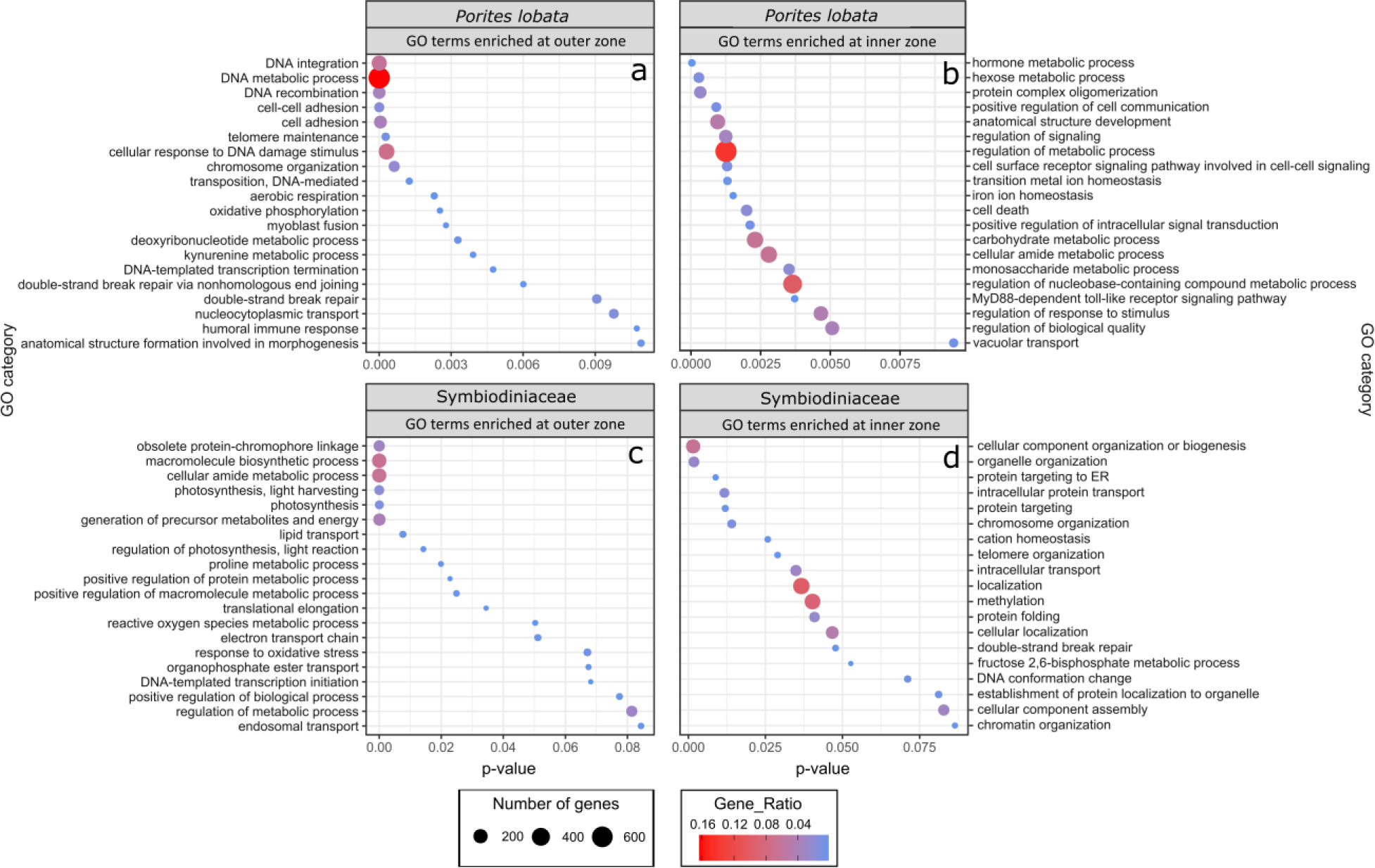
GO terms identified as enriched when comparing inner and outer zone for both *Porites lobata* and Symbiodiniaceae. Up and down regulated genes were separated into “enriched in the Inner zone” and “enriched in the Outer zone”. Gene ratio (color of circles) indicates the number of genes represented by a GO term relative to the total number of differentially expressed genes whereas number of genes (size of circle) is the total number in that GO category.

GO terms associated with aerobic energy generation for *Porites lobata* were enriched in the outer zone compared to the inner zone (Fig. 4a), further indicating that corals in the inner zone experienced hypoxic stress, as can be expected under conditions of severe sedimentation (Weber *et al*., 2012; Bollati *et al*., 2021). In *ex situ* sedimentation experiments, Bollati et al. (2021) found genes associated with an anaerobic glycolytic pathway upregulated in two coral species. Similarly, we find glyceraldehyde-3-phosphate dehydrogenase (GAPDH) (Table S6) upregulated within Symbiodiniaceae in the inner zone and GO terms for aerobic respiration (GO:0009060) and oxidative phosphorylation (GO:0006119) of *Porites lobata* enriched in the outer zone (Fig. 4a), pointing to a reduction of aerobic respiration in corals of the inner zone to mitigate oxygen limitations caused by severe sedimentation. Furthermore, we found a hypoxia-inducible factor (Aryl hydrocarbon receptor nuclear translocator; Table S6) upregulated in *Porites lobata* colonies from the inner zone. While not intuitive at first glance, the observed enrichment of the GO terms response to oxidative stress (GO:0006979) and reactive oxygen species metabolic process (GO:0072593) in Symbiodiniaceae of the outer zone (Fig. 4b) is consistent with metabolic differences between inner and outer zones. In particular, increased photosynthetic activity in the less turbid outer zone compared to the inner zone likely elevated photosynthetic rates and increased ROS production of Symbiodiniaceae that required mitigation. Similarly, antioxidant-associated genes (HSP-70, Thioredoxin domain-containing proteins, Superoxide dismutase, Dnaj-like proteins, peroxidase; Table S6) were upregulated in *Porites lobata* in the outer zone. Our results are consistent with a switch from aerobic to anaerobic-dominated respiration in *Porites lobata* from outer to inner zone. Quantifying the impacts of such metabolic switching on the health of *Porites lobata* using appropriate assays (Murphy and Richmond, 2016) would provide additional insights into the adaptive capacity of this resilient coral species to chronic sedimentation stress and resulting hypoxia.

### Immune response

Persisting under severe sedimentation that arrives in pulses associated with rainfall events requires rapid sensing of environmental change and signal transduction to initiate metabolic responses. We found GO terms associated with positive regulation of cell communication (GO:0007267), positive regulation of signal transduction (GO:0009967), MyD88-dependent toll-like receptor signaling pathway (GO:0002755), cell surface receptor signaling pathway involved in cell-cell signaling (GO:0007166), and regulation of response to stimulus (GO:0048583) overrepresented in *Porites lobata* growing in the inner zone impacted by severe sedimentation (Fig. 4b). The MyD88-dependent toll-like receptor signaling pathway is involved in the innate immune response of corals (Mydlarz *et al*., 2016), stimulating the expression of pro-inflammatory cytokines (Poole & Weis, 2014). In sponges, this signaling pathway is strongly upregulated in response to bacterial endotoxin lipopolysaccharide, effectively eliminating gram-negative bacteria through the formation of a recombinant protein (Wiens *et al*., 2005). Gram-negative pathogenic bacteria, such as *Vibrio* spp., are associated with marine sediment (Franco *et al*., 2020) and bacterial communities associated with heavy sediment loads are known exacerbate mortality in corals (Hodgson, 1990; Weber *et al*., 2012). Indeed, microbiome metabarcoding previously revealed that *Porites lobata* colonies growing in the inner zone of Fouha Bay, close to the river mouth, harbored higher relative abundances of Vibrionaceae than colonies in the outer zone (Fifer *et al*., 2022).

Although corals are thought to only possess an innate immune system (Palmer and Traylor-Knowles, 2012), presence of the MyD88-dependent toll-like receptor signaling pathway is an example of an adaptive-like immune response system that may help corals mitigate bacterial infections (Wiens *et al*., 2005; Poole and Weis, 2014) associated with severe sedimentation. Only recently have coral immune cells and their gene expression been characterized using single cell transcriptomics (Levy *et al*., 2021). Interestingly, we found genes associated with immune functions in coral immune cells upregulated in the inner zone (Interferon regulatory factor 2, Tyrosinase, and Homeobox genes coding for Meis2 and Nkx-2.2a; Table S6). In summary, we saw enrichment of cell signaling GO terms (Fig. 4b) and upregulation of genes previously identified as molecular signatures of immune function (Palmer and Traylor-Knowles, 2012; Poole and Weis, 2014; Mydlarz *et al*., 2016; Levy *et al*., 2021) in *Porites lobata* colonies growing in the inner, severely sedimentation-impacted zone, suggesting that coral colonies in the inner zone rely on their immune response to mitigate the impacts of severe sedimentation.

Despite environmental sensing and resulting metabolic and immune responses, cell damage under severe stress is expected to increase. Initiation of apoptosis is a critical process that allows organisms to maintain tissue homeostasis by eliminating damaged cells that, for example, pose a risk for infection by pathogens (Ameisen, 2002). Various programmed cell death pathways have been identified in the cnidarian stress response, but destabilization of cellular adhesion seems to be a crucial component in apoptosis initiation, given that changes in cell adhesion proteins have been associated with the response to coral bleaching (Gates, Baghdasarian and Muscatine, 1992; DeSalvo *et al*., 2012; Traylor-Knowles and Connelly, 2017), disease (Daniels *et al*., 2015), and heat plus sedimentation stress (Poquita-Du *et al*., 2019).

Consistent with these prior studies, we found GO terms for cell death (GO:0008219) enriched in the inner zone (Fig. 4b) as well as enrichment of cell adhesion terms (GO:0007155, GO:0098609) in the outer zone (Fig. 4a). This pattern suggests that *Porites lobata* from the inner zone modified the extracellular matrix of part of its population of cells in conjunction with elevated rates of apoptosis initiation to remove cells damaged by severe sedimentation stress in an attempt to survive in this turbid, marginal habitat.

### Conclusions

We provide a framework of the metabolic acclimation response of *Porites lobata* and its Symbiodiniaceae endosymbionts to sedimentation stress under *in situ* field conditions. We identified processes that may allow for sedimentation tolerant corals, such as *Porites lobata*, to persist in habitats impacted by chronic and severe sedimentation. Switching between energy generation pathways may help corals living in sedimentation-impacted, turbid reefs to maintain a stress response to survive while light and oxygen availability are diminished. Rapid environmental sensing and cell-cell communication allows for corals to respond to freshwater influx, sedimentation stress, and bacterial challenges associated with terrestrial runoff. Given the cellular damage associated with environmental stress, removal of damaged cells through programmed cell death pathways is crucial to maintain coral colony integrity and close entryways to pathogens. These putative mechanisms identified by us using transcriptomics ought to be further investigated and tested using additional approaches from the multi-omics toolbox under field and controlled conditions to gain a better understanding of sedimentation stress tolerance mechanisms in corals. Coastal development activities are continuously increasing sedimentation and turbidity in near-shore reefs and a more in-depth understanding of the metabolic plasticity of diverse corals will facilitate predicting the acclimation and adaptation limits of near-shore reef ecosystems in the Anthropocene.

## Acknowledgements

This work was supported through National Science Foundation EPSCoR awards OIA-1457769 and OIA-1946352. We would like to thank Constance Sartor for assisting with genetics lab work and Edriel Aquino for processing sediment samples.

## Collection permissions

Coral and sediment samples were collected under University of Guam Marine Laboratory collection permits issued by Guam’s Division of Aquatic & Wildlife Resources, complying with local laws and regulations.

## Author contributions

MMG and BB conceived of the project. MMG conducted field work and collected samples and environmental data. MMG and CL performed genetics laboratory work. CL and BB analyzed the genetic data. CL drafted the manuscript; all authors reviewed and revised the manuscript.

## Data availability

Sequencing data and transcriptome assemblies were deposited in NCBI’s GenBank under BioProject <u>PRJNA914839</u>. Genebank accession numbers for coral host data are found in figure 3 and in the supplementary tables. Results of differential gene expression and GO enrichment analyses are provided in the supplementary tables accompanying this article.

